# The role of equilibrium cost in the evolution of honest signalling: waste or optimal investment?

**DOI:** 10.1101/256370

**Authors:** Szabolcs Számadó, Dustin J. Penn

## Abstract

The relationship between signal cost and honesty is a controversial and unresolved issue. The handicap principle assumes that signals must be costly at equilibrium to be honest, and the greater the cost, the more reliable the signal. However, theoretical models and simulations question the necessity of equilibrium cost for the evolution of honest signalling. Honest signals can evolve without costs, and they can evolve through differential benefits with no need for differential costs. Here we investigate the role of equilibrium signal cost in the evolution of honest signals in both differential benefit and differential cost models using an agent-based simulation. We found that there is an optimal investment paid by honest individual that allows for the highest level of honesty when there is correlation between signal cost paid by low and high-quality individuals. This holds for both differential benefit and differential cost models as long there is a correlation between signal cost paid by low and high quality individuals. However, increasing equilibrium signal cost poses an obstacle and hinders the evolution of honest signalling when there is no correlation between the cost paid by low and high-quality individuals. Last but not least, we found that the potential cost of cheating is a much better predictor of honesty than the equilibrium cost paid by honest signallers.

## 1. Introduction

The evolution of honest signalling remains a controversial problem. The most cited theory is the so-called handicap principle [1-3]. There are several versions of this hypothesis, which all assume that signals must be costly at the evolutionary equilibrium to be honest, and that the honest signals evolve because rather than despite of their costs. The most-cited version concludes that honest signals of individual quality evolve when low quality individuals pay higher costs for signalling compared to high quality individuals (strategic or *differential costs* handicap model) [2]. Zahavi argued that honest signals evolve through a special type of selection, called ‘signal selection’ that mysteriously favours waste rather than efficiency [3]. Yet, there have been an increasing number of theoretical studies that challenge the claim that signals must be costly at equilibrium to be honest [4-10].

The evolution of honest signals has been investigated in several game theoretical models. These signalling games have shown the existence of various solutions from non-signalling [11], through pooling [12] to costly [2, 13, 14] or cheap separating equilibria [4-6]. Multiple equilibria often coexist in a single model [6, 15-17]. Unfortunately, standard game theory cannot predict which one of these equilibria will be reached by evolution [17]. An evolutionary approach is needed to resolve this problem, which can make predictions on the trajectory of evolving populations.

Individual or agent-based models provide an alternative approach to study such evolutionary trajectories [11, 18-22]. These models have highlighted different aspects of selection for honest signals, such as the evolution (or lack of) of honest and cheating strategies in a game of aggressive communication [18-20], the instability of costly equilibria in parent-offspring communication [11] or the role of signal competition in the evolution of costly signals [22]. Recently, Kane & Zollman [21] investigated the different attractors of honest costly versus hybrid equilibria in a differential benefit model of a simple action-response game. They found that the hybrid equilibrium evolved more frequently. They also found that low cost of both signals favours the evolution of costly honest equilibria. In other words, costly equilibria evolve but only if it is not too costly [21].

While the above models investigate interesting problems none of them investigates systematically the role of equilibrium on honesty in both differential benefit and differential cost models. Here we this question. Our goal is to determine the relationship between the signal cost paid by honest signallers at the equilibrium and the honesty of the system. We considered two conditions: (i) when there is no correlation between the signal cost for low and high-quality individuals (differential cost model), and (ii) with correlation between the signal cost for low and high-quality individuals (differential benefit and differential cost model). We modelled the evolution of signalling by means of individual based simulations. We found that honesty evolved more readily at intermediate values of equilibrium signal cost when there is a correlation between signal cost paid by low and high-quality individuals, but there is a monotone decreasing relation between honesty and equilibrium signal cost when there is no such correlation.

## 2. The model

We used a simple signalling game known as an action-response game [4, 5, 14, 23]. This type of signalling game is used to describe situations in which receivers control a (non-divisible) resource that signallers wish to obtain, such as during mate assessment or parent-offspring conflict [24]. There is a *signaller* and a *receiver*; the receiver holds an indivisible resource. Signallers can be either high or low quality, and their quality is not known and cannot be observed by the receivers. Signallers are always better off with the resource, and receivers are only interested in transferring the resource to high quality signallers. The signallers can opt to display a signal, which may or may not be an honest signal of quality.

The receivers’ fitness (*F*_*r*_) depends both on the signaller’s quality (*a*), which can be high (*H*) or low (*L*) and on the receiver’s response (*z*), which can be up (*U*): to give the resource, or down (*D*): not to give the resource. The signaller’s fitness (*F*_*s*_) is the sum of the value of the receiver’s response (*V*), minus the cost of signalling (*C*). The value of the receiver’s response (*V*) both depends on the quality of the signaller and on the receiver’s response; while the cost of signals (*C*) can depends on both the quality of the signaller and on the signaller’s behaviour (*b*_*s*_), which can be to signal (*S*) or not to signal (*N*). The receivers’ (*F*_*r*_) and the signallers’ (*F_s_*) fitness can be written up as follows:

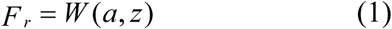

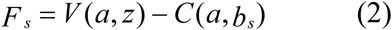

The fitness of both the receiver and the signaller can depend on the survival of the other player (*r*); in the simplest case this could imply that they are relatives; alternatively, they can help each other in some other way or they can belong to the same group (see Maynard Smith, 1991). With the help of *r* it is both possible to describe situations where this interdependence is high (*r*>>0, for example parent-offspring communication) or situation where there is no relatedness and the players do not interact with each other outside the signalling game (i.e. *r*=0). Based on these assumptions the inclusive fitness of the signaller (*E*_*s*_) and the receiver (*E*_*r*_) can be written as follows:

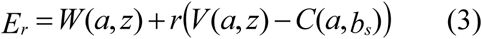

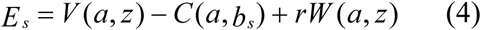

Let *V*_*h*_ and *V*_*l*_ denote the difference in fitness for high-, and low-quality signaller respectively between obtaining the resource or not [4, 5]:

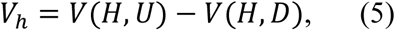

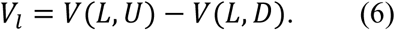

We can define *W*_*h*_, *W*_*l*_ and *C*_*h*_, *C*_*l*_ in a similar way:

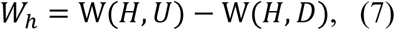

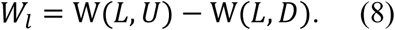

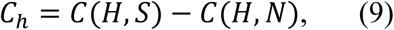

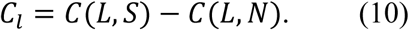

This notation will be used in the rest of the article (see Table 1. for a summary). Figure 1 depicts the signalling game and Table 2 describes the pay-offs.

**Figure 1.**
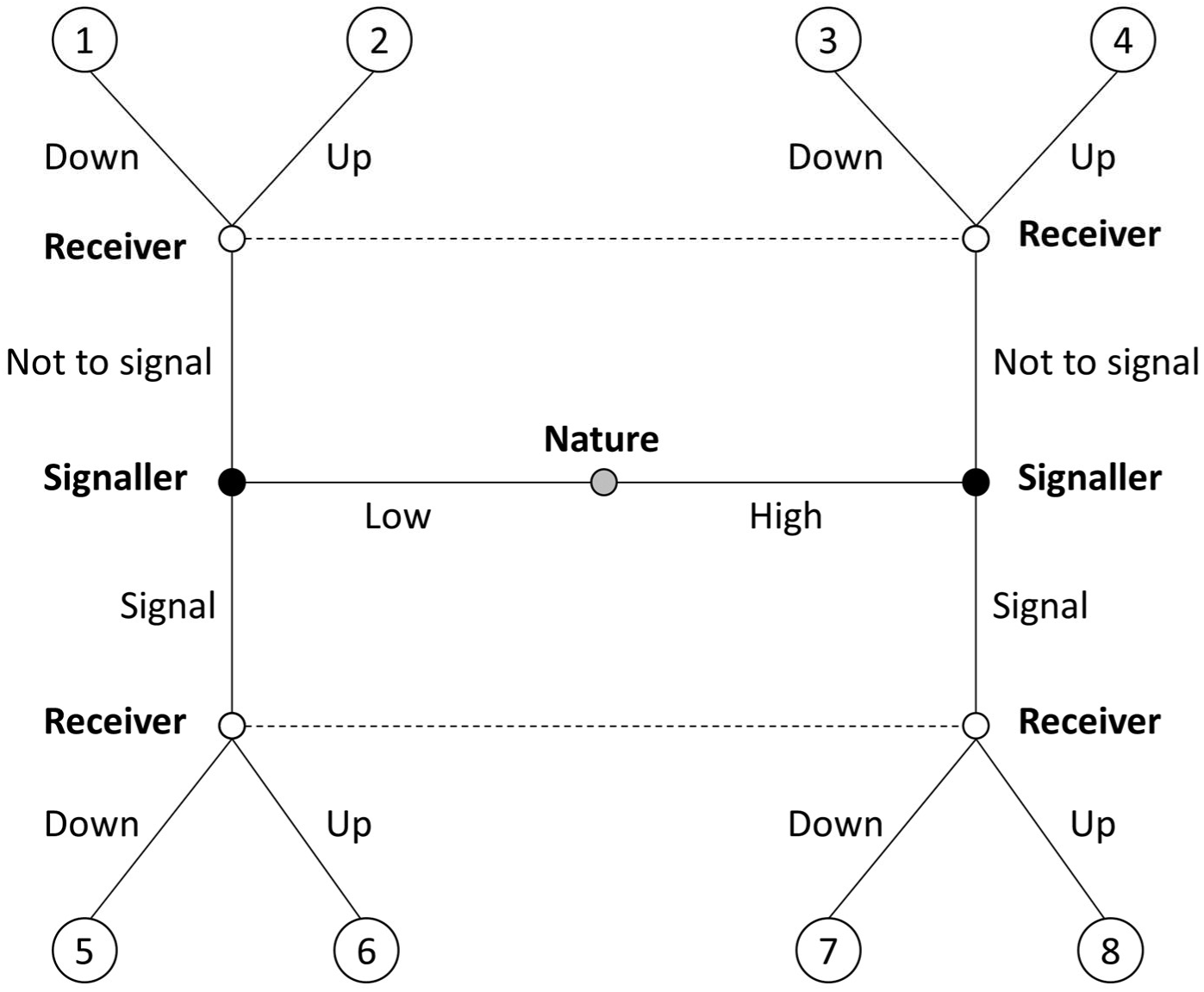
The action-response game. Nature first decides the type of the signaller, signallers can decide to signal or not to signal, and finally receivers can decide to transfer the resource to the signaller (Up) or not to do it (Down).

**Table 1.**
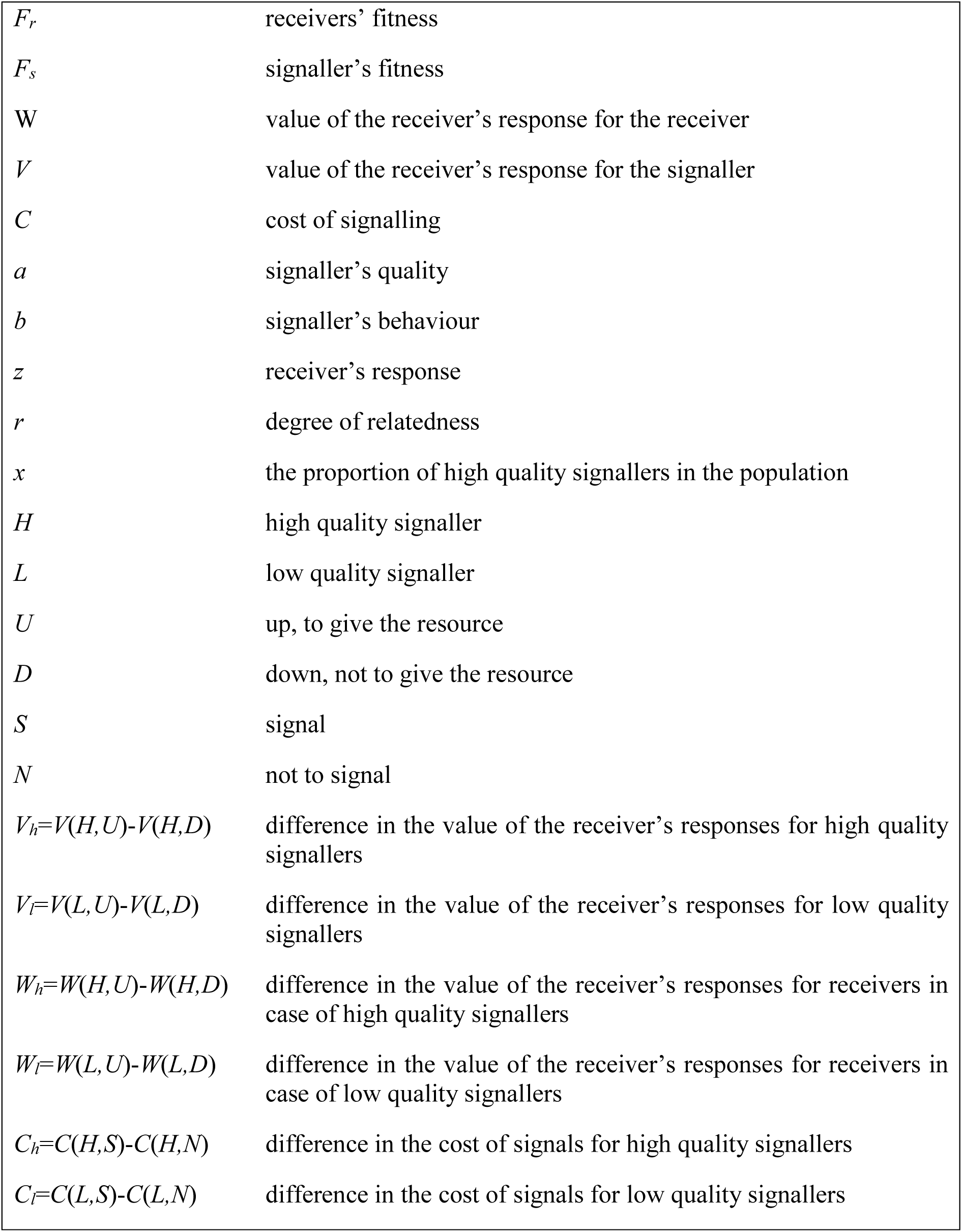
Parameters and the notation of the model.

**Table 2.** Pay-offs of the model. See the end nodes in Figure 1.

| Receiver's fitness |  | Signaller's fitness | Receiver's fitness | Signaller's fitness |
| --- | --- | --- | --- | --- |
| **1,** | $L_r = W(L, D) + rV(L, D)$ | $L_s = V(L, D) + rW(L, D) - C(L, N)$ | 1 | 1 |
| **2,** | $L_r = W(L, U) + rV(L, U)$ | $L_s = V(L, U) + rW(L, U) - C(L, U)$ | 0 | 2 - dV |
| **3,** | $L_r = W(H, D) + rV(H, D)$ | $L_s = V(H, D) + rW(H, D) - C(H, N)$ | 0 | 1 |
| **4,** | $L_r = W(H, U) + rV(H, U)$ | $L_s = V(H, U) + rW(H, U) - C(H, N)$ | 1 | 2 - dV |
| **5,** | $L_r = W(L, D) + rV(L, D)$ | $L_s = V(L, D) + rW(L, D) - C(L, S)$ | 1 | 1 - C(H, S) - dC |
| **6,** | $L_r = W(L, U) + rV(L, U)$ | $L_s = V(L, U) + rW(L, U) - C(L, S)$ | 0 | 2 - C(H, S) - dC |
| **7,** | $L_r = W(H, D) + rV(H, D)$ | $L_s = V(H, D) + rW(H, D) - C(H, S)$ | 0 | 1 - C(H, S) |
| **8,** | $L_r = W(H, U) + rV(H, U)$ | $L_s = V(H, U) + rW(H, U) - C(H, S)$ | 1 | 2 - C(H, S) |

The last step is to identify the available strategies for receivers and signallers. Signallers must decide their move based on the state Nature assigned to them, whereas receivers have to decide based on the signal they receive from signallers. Let *Q*_*S*_(*, *) denote signaller’s strategy where the first positions tells what to do if Low state and the second tells what to do if High. Let’s *Q*_*R*_(*, *) denote receiver’s strategy where the first positions tells what to do if the signallers sent No signal and the second tells what to do if the signaller sent Signal. For example, *Q*_*S*_(S, N) denotes a pure strategy where individuals Signal if in Low state and they give No signal if they are in High state. Whereas, for example, *Q*_*R*_(U, U) denotes a receiver strategy that gives the resource to the signaller regardless of the signal.

There are four types of honest signalling equilibria in this game, where signallers give a variable signal and receivers response selectively to it. These four types can be analysed in the same way. Here we focus on the “traditional” signalling equilibrium where only high-quality signallers give signal and receivers give the resource only to those signallers who signal. This equilibrium is defined by the following signaller and receiver strategy pair: *Q*_*S*_(N, S) and *Q*_*R*_(D, U).

## 3. Methods

We used individual based computer simulations to assess the evolutionary trajectories of different populations. The computer simulations model a population of individuals (*n*=100) playing an action-response game. The structure of the game corresponds to the analytical model (see Figure 1). Signaller and receiver behaviour coded by “genes” that can be inherited into the next generation. Both signallers and receiver have two genes. The first gene of signallers specifies which signal to give when low quality, the second one specifies which signal to give when high quality. The first gene of receivers specifies which response to give when the signaller did not send a signal, the second specifies which response to give in case of a signal. Each scenario was seeded with a random mix of signaller and receiver strategies.

Each individual plays ten games against randomly chosen opponents in one step of the simulations. Reproductive success is proportional to the pay-offs received when playing this action-response game. Individuals produced offspring proportional to their fitness. Out of the pool of offspring one offspring was selected randomly; this offspring replaced a randomly selected individual of the original population. The offspring inherits its genes from its parent with a mutation probability =0.05. Simulations were iterated for *i*=20000 steps. For each parameter combination m=100 independent runs were implemented. The last step of these runs was taken and analysed in detail. The timeline of each run was also saved and analysed.

We manipulated the comparative advantage of high quality signallers in four different scenarios. We changed the benefit difference between high and low-quality signallers in the differential benefit models (d*V*), or we changed the cost difference between low and high quality signallers in the differential cost scenarios (d*C*). We changed the cost of signalling for high quality signallers (*C*_*h*_) from zero to one with steps 0.1. The cost of signal for low quality individuals (*C*_*l*_) was either the same in differential benefit models, or it was the cost of signal for high quality individuals plus the cost difference in the scenario (d*C*). These scenarios and the corresponding parameters are summed up in Table 3 and 4.

**Table 3.** Correlated cost.

|  | differential<br>benefit models |  | differential<br>cost models |  |  |  | Expected<br>equilibrium |
| --- | --- | --- | --- | --- | --- | --- | --- |
| | $dV$ | $dC$ | $dV$ | $dC$ | $C(H,S)$ | $C(L,S)$ | |
| 1. | 0.3 | 0 | 0 | 0.3 | (from 0 to 1, steps 0.1) | $C(H,S) + dC$ | Hybrid |
| 2. | 0.5 | 0 | 0 | 0.5 | (from 0 to 1, steps 0.1) | $C(H,S) + dC$ | Hybrid |
| 3. | 0.7 | 0 | 0 | 0.7 | (from 0 to 1, steps 0.1) | $C(H,S) + dC$ | Hybrid |
| 4. | 0.9 | 0 | 0 | 0.9 | (from 0 to 1, steps 0.1) | $C(H,S) + dC$ | Hybrid |
| 5. | 1.1 | 0 | 0 | 1.1 | (from 0 to 1, steps 0.1) | $C(H,S) + dC$ | Honest |
| 6. | 1.3 | 0 | 0 | 1.3 | (from 0 to 1, steps 0.1) | $C(H,S) + dC$ | Honest |
| 7. | 1.5 | 0 | 0 | 1.5 | (from 0 to 1, steps 0.1) | $C(H,S) + dC$ | Honest |
| 8. | 1.7 | 0 | 0 | 1.7 | (from 0 to 1, steps 0.1) | $C(H,S) + dC$ | Honest |

**Table 4.**
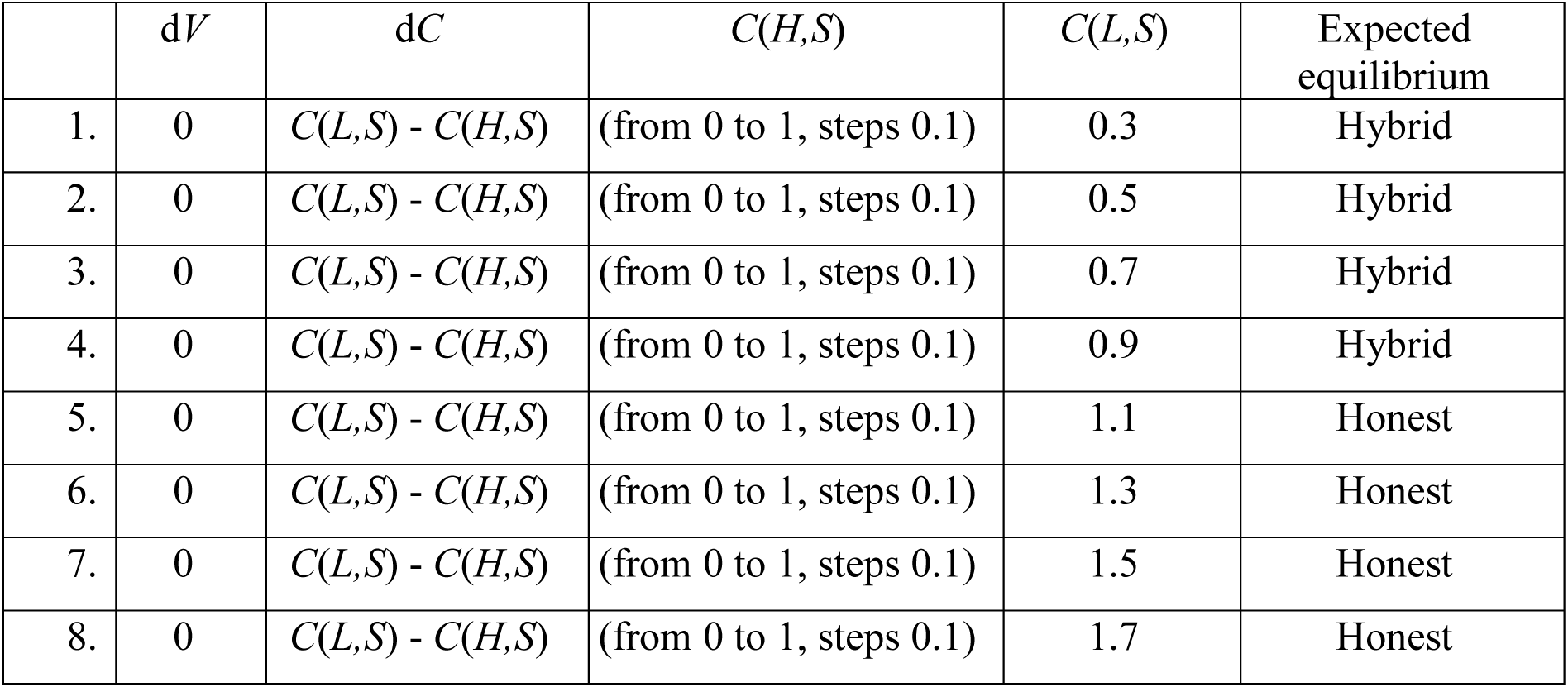
Differential cost models, uncorrelated cost.

| | $dV$ | $dC$ | $C(H,S)$ | $C(L,S)$ | Expected<br>equilibrium |
| --- | --- | --- | --- | --- | --- |
| 1. | 0 | $C(L,S) - C(H,S)$ | (from 0 to 1, steps 0.1) | 0.3 | Hybrid |
| 2. | 0 | $C(L,S) - C(H,S)$ | (from 0 to 1, steps 0.1) | 0.5 | Hybrid |
| 3. | 0 | $C(L,S) - C(H,S)$ | (from 0 to 1, steps 0.1) | 0.7 | Hybrid |
| 4. | 0 | $C(L,S) - C(H,S)$ | (from 0 to 1, steps 0.1) | 0.9 | Hybrid |
| 5. | 0 | $C(L,S) - C(H,S)$ | (from 0 to 1, steps 0.1) | 1.1 | Honest |
| 6. | 0 | $C(L,S) - C(H,S)$ | (from 0 to 1, steps 0.1) | 1.3 | Honest |
| 7. | 0 | $C(L,S) - C(H,S)$ | (from 0 to 1, steps 0.1) | 1.5 | Honest |
| 8. | 0 | $C(L,S) - C(H,S)$ | (from 0 to 1, steps 0.1) | 1.7 | Honest |

We measured several properties of the system including honesty, trust and the potential cost of cheating. This potential cost can be defined as the cost for low quality individuals to mimic the intensity of signals used by high quality individuals:

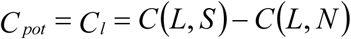

Honesty (*H*) measures the correlation between signal and the underlying quality [8](Számadó, 2011), and thus it can vary between-1 and 1. Trust can vary between 0 and 1 and it measures how willing are receivers to give the resource to signallers who give signals higher than the lowest signal intensity.

## 4. Results

There is a clear optimal investment for equilibrium signal cost in both differential benefit and differential cost models with correlated signal cost. Differential cost models perform better and achieve higher levels of honesty than differential benefit ones (see Figure 2). Figure 3 shows a more detailed breakdown of these results. It is clear that there is an optimum value of signal cost for high quality signallers in differential benefit models regardless of the region (hybrid vs. honest; see Fig 3.a vs 3.c). On the other hand, differential cost models have clear optimum only in the hybrid region (Fig 3.b); increasing the equilibrium cost of signalling has no effect on honesty in the honest region up to a threshold (the cost for high quality individuals, *C*_*h*_=0.5; see Fig 3.d). After this threshold, honesty declines rapidly with increasing signal cost for high quality signallers.

**Figure 2.**
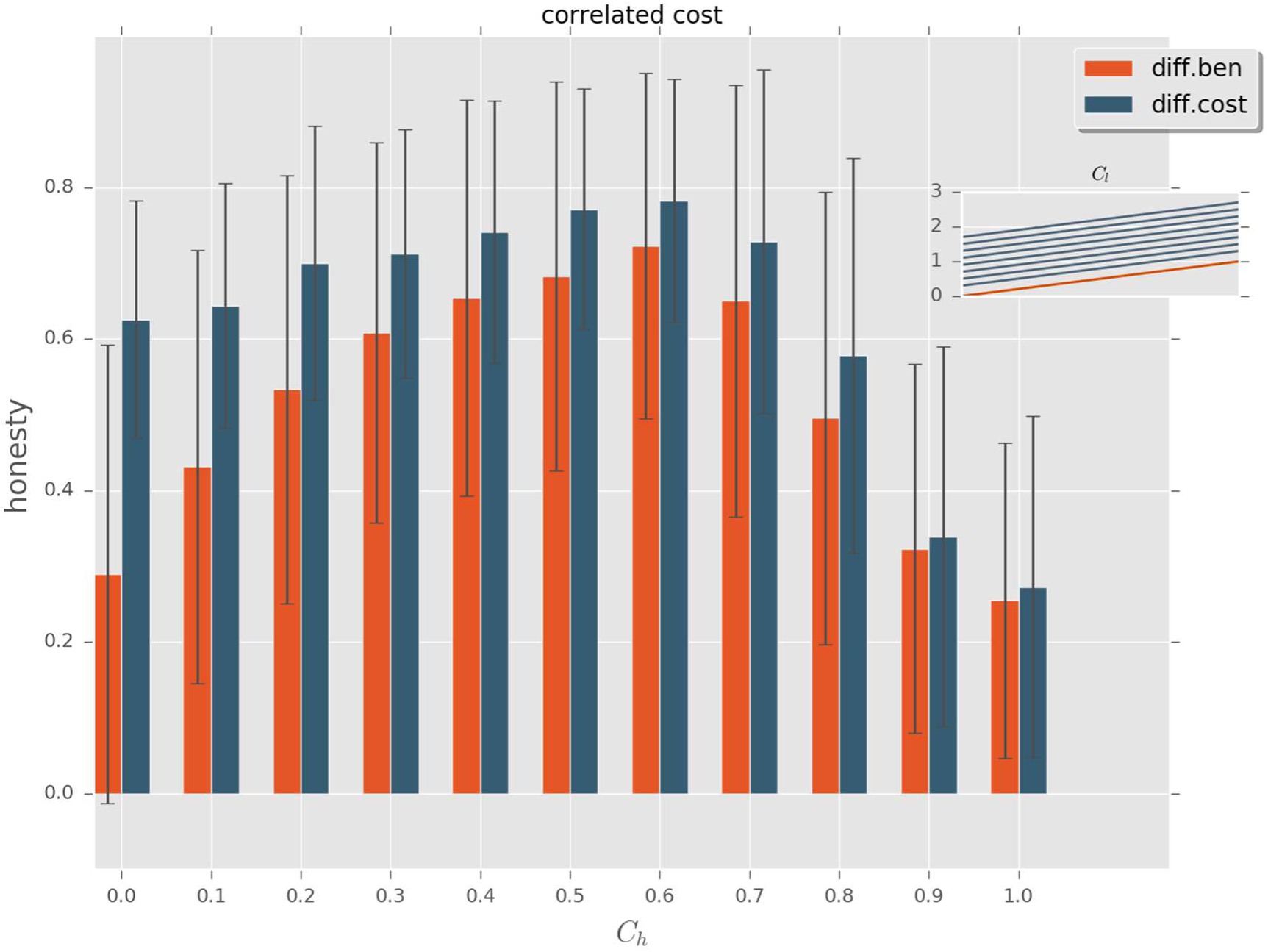
The average level of honesty as function of signal cost for high quality individuals (*C*_*h*_); correlated cost. There is a correlation between the cost paid by low and high-quality individuals. The small inlet shows the change of cost for low quality individuals as function of *C*_*h*_. Red: differential benefit models; blue: differential cost models.

**Figure 3.**
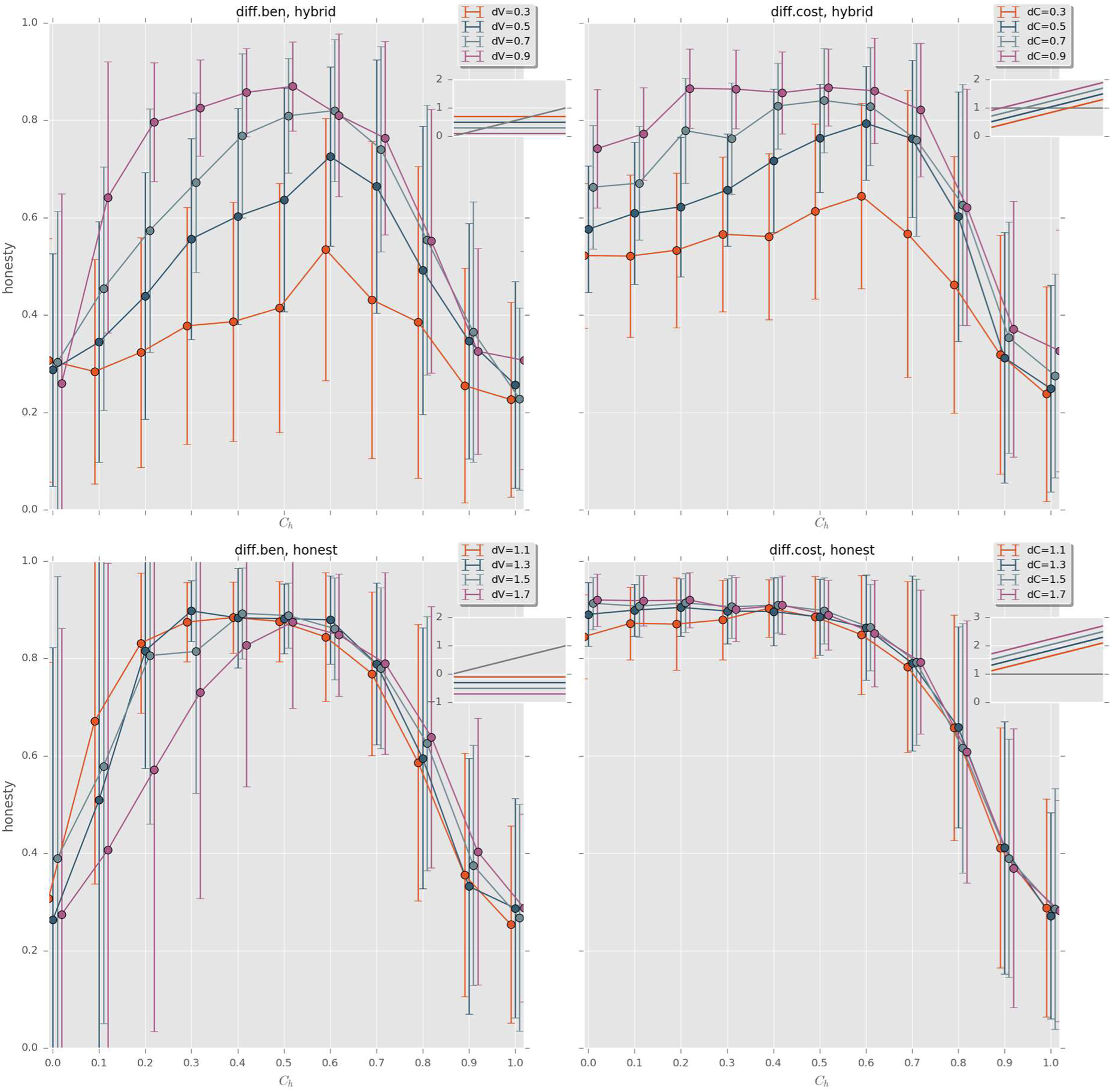
The average level of honesty as function of signal cost for high quality individuals (*C*_*h*_) in four different scenarios. (a) Differential benefit models with different comparative advantage (d*V* ranging from 0,3-0,9; hybrid region); (b) differential cost models with different comparative advantage (d*C* ranging from 0,3-0,9; hybrid region); (c) differential benefit models with different comparative advantage (d*V* ranging from 1,1-1,7; honest region); (d) differential cost models with different comparative advantage (d*C* ranging from 1,1-1,7; honest region). Small inlet shows the change of cost for low quality individuals (*C*_*l*_) as function of *C*_*h*_ in each of these scenarios.

Increasing the signal cost for high quality signallers is clearly an obstacle for the evolution of honesty in the hybrid region in the case of uncorrelated cost (Fig 4.). Honesty seems to be robust against the increase of costs for high quality signallers (*C*_*h*_) in the honest region up to some point, just like before (Fig 3.d), and afterwards honesty rapidly declines with increasing *C*_*h*_. All in all, there is no region in the uncorrelated cost model where increasing signal cost for honest signallers would promote the evolution of honest signalling.

**Figure 4.**
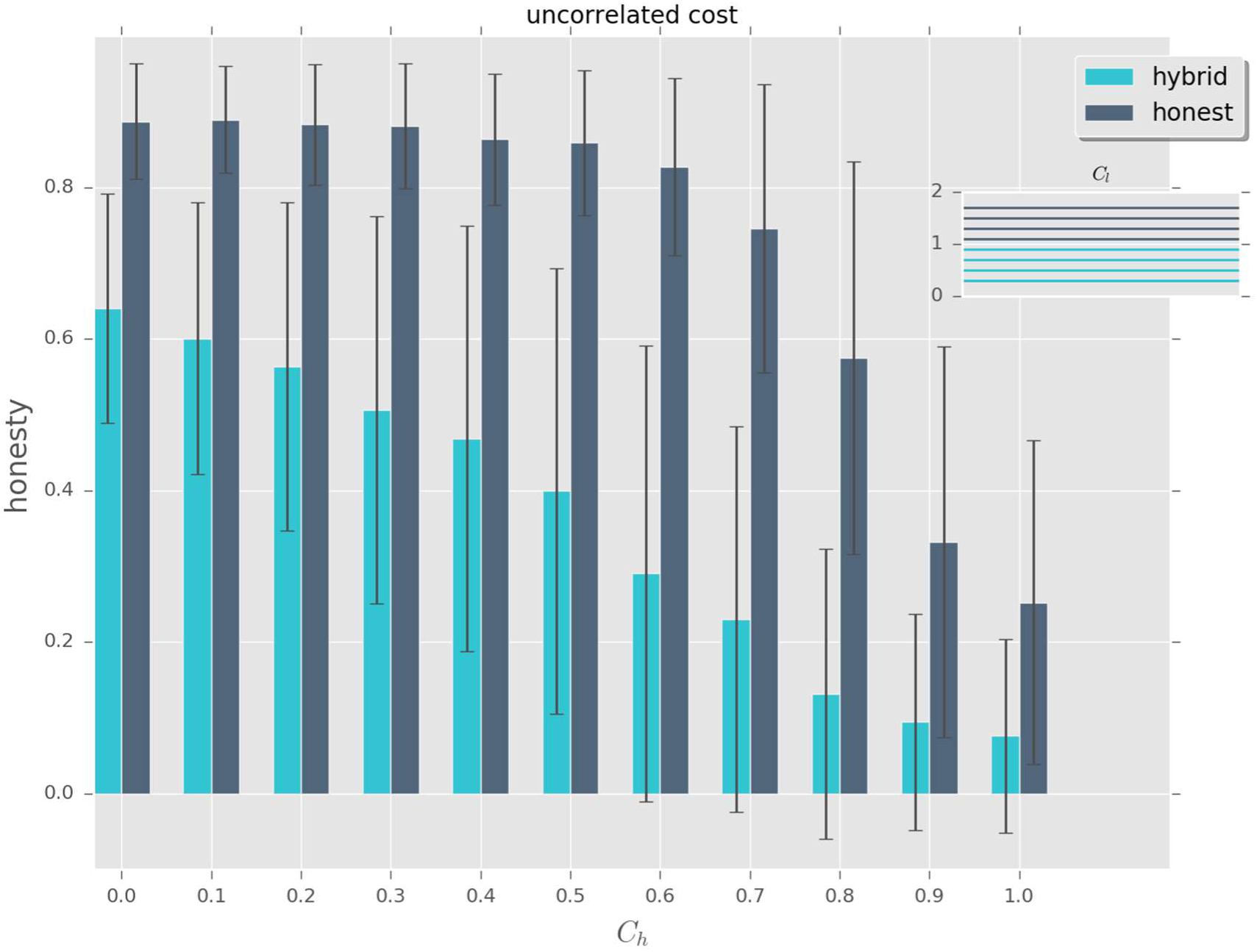
The average level of honesty as function of signal cost for high quality individuals (*C*_*h*_); uncorrelated cost. There is no correlation between the cost paid by low and high-quality individuals. The small inlet shows the change of cost for low quality individuals as function of *C*_*h*_. Light blue: hybrid regions; blue: honest regions.

We have also made “heat maps” to show the trajectories of individual runs in the trust-honesty state-space. Populations show random walk like behaviour at low potential cost of cheating (Fig 5,6), whereas populations quickly converge into the high trust-high honesty region with high potential cost of cheating (Fig 7,8). Appendix 1 shows the individual timelines of the runs shown in Figs 5-8.

**Figure 5.**
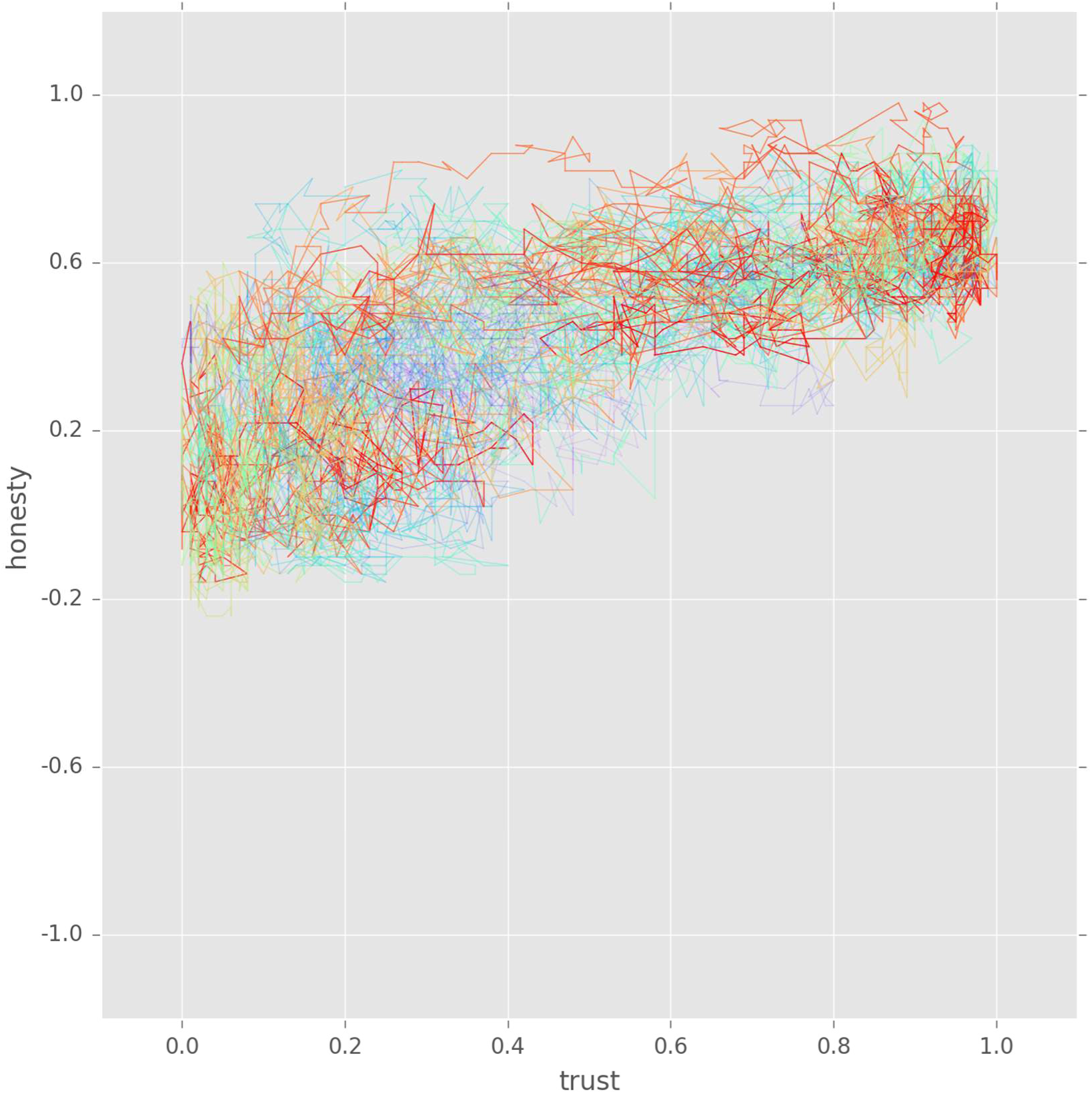
Heat-map. Trajectories of 10 populations in the trust-honesty state-space. From blue to red: from early steps to last steps. Differential benefit model, parameters: d*V*=0.5, *C*=0.2.

**Figure 6.**
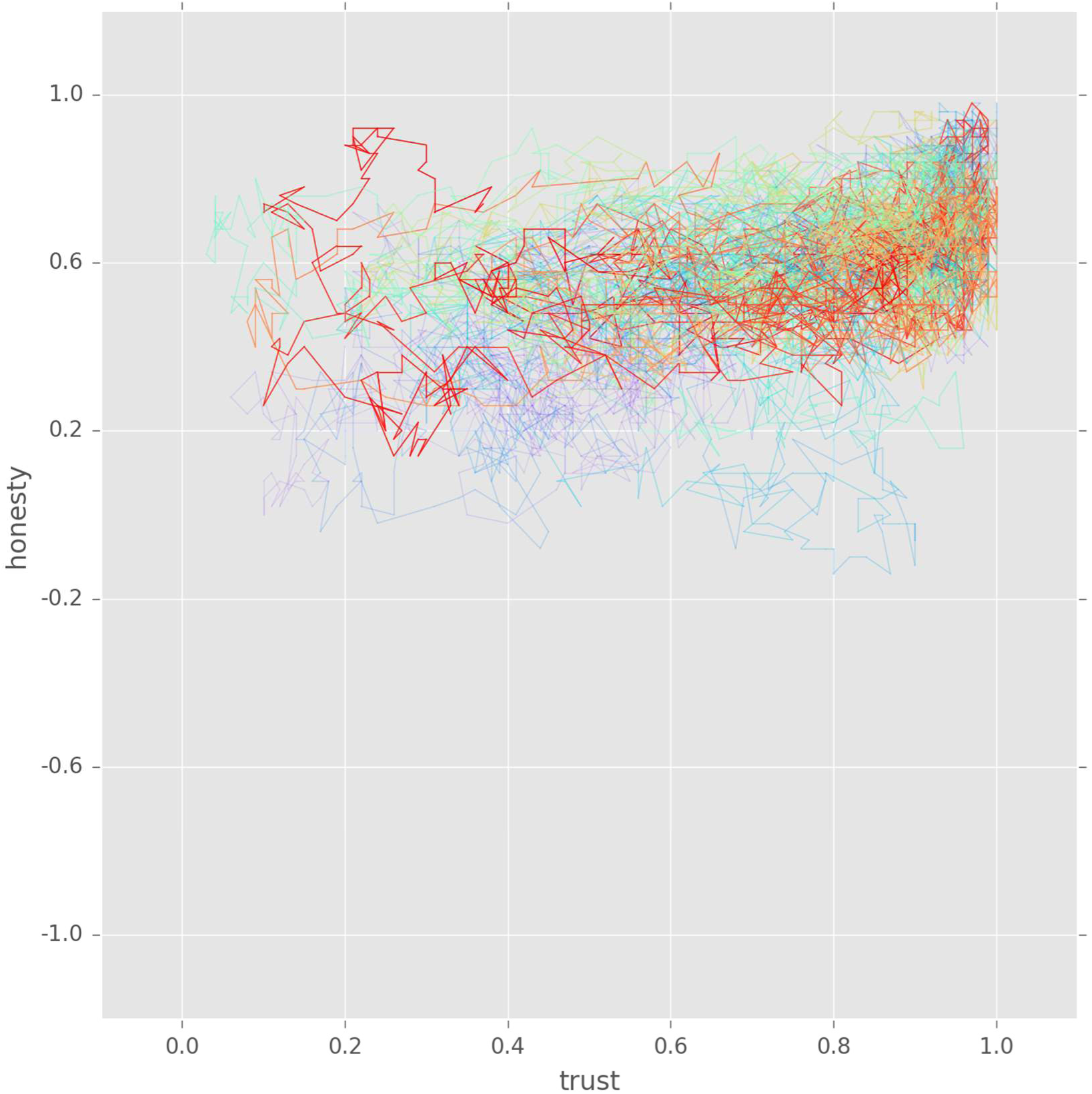
Heat-map. Trajectories of 10 populations in the trust-honesty state-space. From blue to red: from early steps to last steps. Differential cost model: d*C*=0.5, *C*_*h*_=0.2.

**Figure 7.**
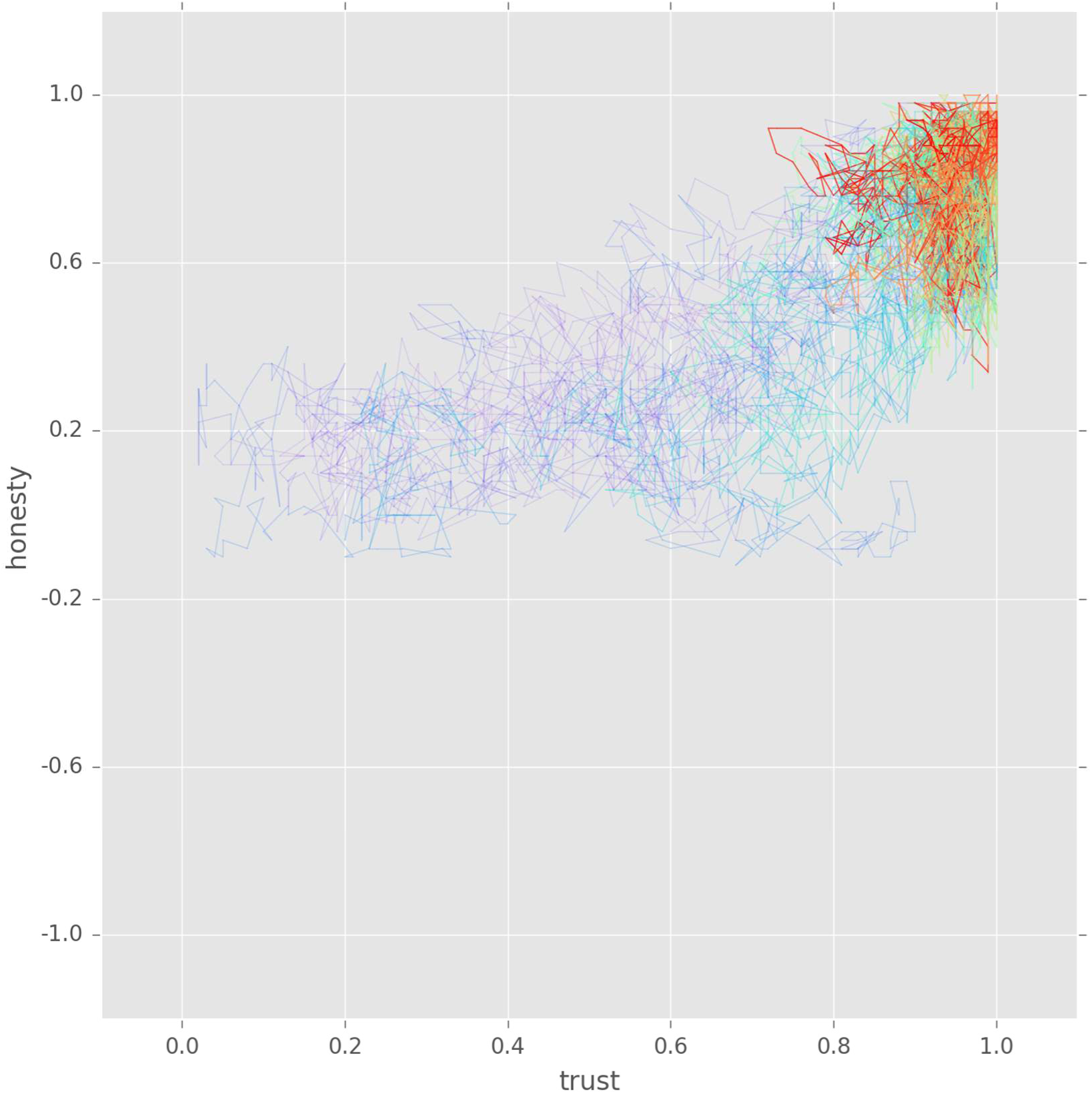
Heat-map. Trajectories of 10 populations in the trust-honesty state-space. From blue to red: from early steps to last steps. Differential benefit model, parameters: d*V*=0.5, *C*=0.6.

**Figure 8.**
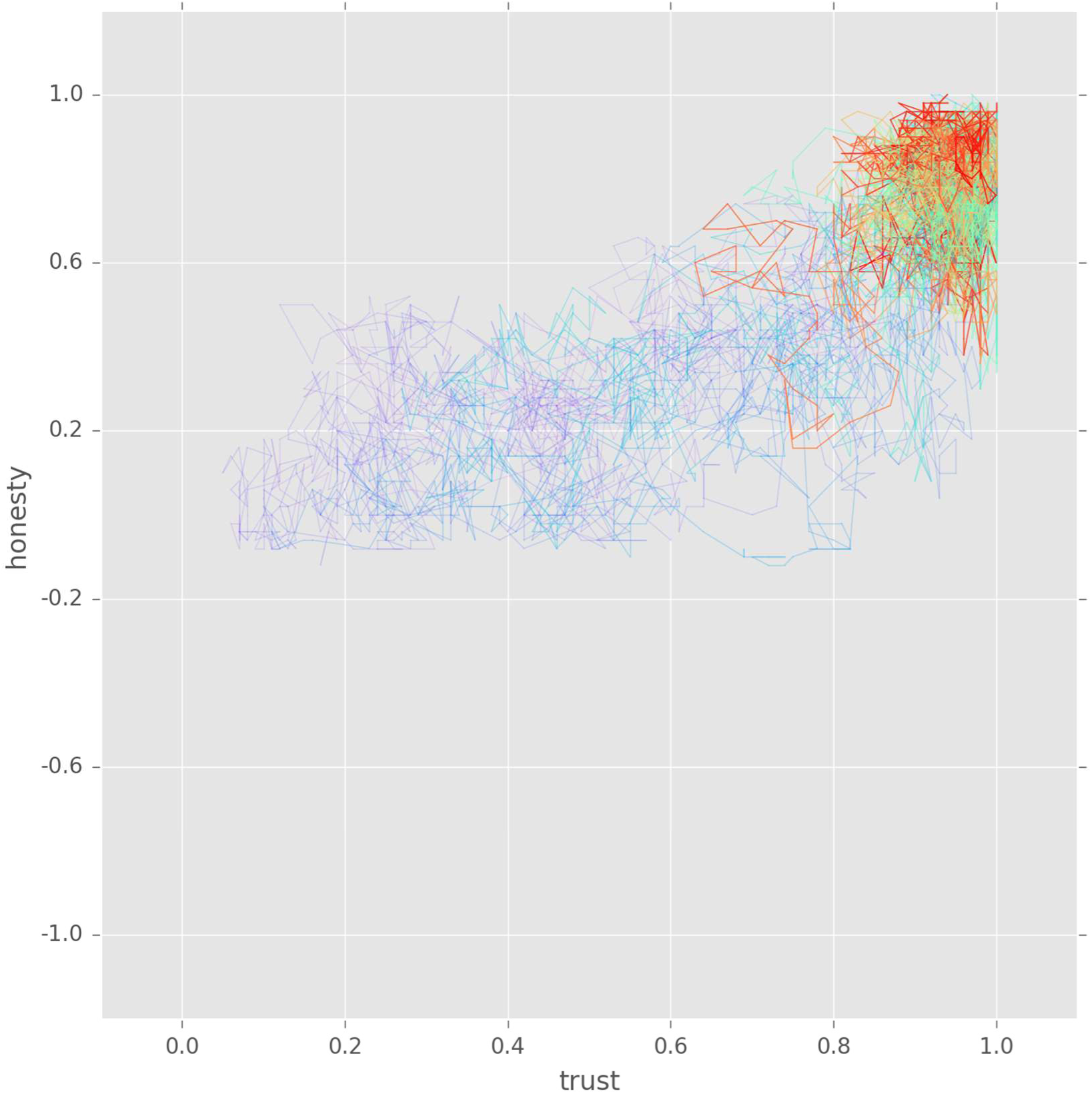
Heat-map. Trajectories of 10 populations in the trust-honesty state-space. From blue to red: from early steps to last steps. Differential cost model: d*C*=0.5, *C*_*h*_=0.6.

## 5. Discussion

Here we studied the co-evolution of signal cost and honesty in a simple action-response signalling game using individual based computer simulations. Our results show that increasing the equilibrium cost for honest (high quality) signallers promotes the evolution of honesty only as long as (i) it contributes to the increase of the *potential cost* of cheating, and (ii) it is not close or higher than the potential benefits of signalling for high quality signallers. These results indicate that there is clear optimum for equilibrium cost both in differential benefit and differential cost models – though only as long as the signal costs of low and high-quality individuals are correlated. If there is no such correlation between these costs, then the increase of equilibrium cost is detrimental to the evolution of honest signalling. Last but not least, the potential cost of cheating is a much better predictor of the evolution of honesty than the equilibrium cost of signals.

Our simulations confirmed several predictions of previous analytical models (see [8] for a review): (i) The cost of producing honest signals at equilibrium in differential cost models can be zero or even negative [4-6]. (ii) Positive equilibrium cost of signals is neither a necessary nor a sufficient condition for honesty [5]. (iii) Finally, honesty is maintained by the potential cost of cheating [4, 5, 8], i.e. honest signalling evolve if the marginal cost is greater than the marginal benefit of cheating [6, 8, 16].

All in all, increasing the equilibrium cost of signals only promotes the evolution of honesty as long as it contributes to the potential cost of cheating (in differential benefit and in correlated cost models); however, even in these scenarios there is an optimum value of equilibrium cost. The realized cost paid by honest signallers actually hinders the evolution of honest signalling when it is uncorrelated with the potential cost of cheating.

These results have important implications for Zahavi’s handicap principle[3]. The predictions of Zahavi can be broadly classified into equilibrium predictions and predictions about the expected evolutionary trajectory of signal evolution. The equilibrium prediction is that selection favours honest signals and that honesty is maintained by a wasteful signal cost.

The evolutionary prediction is that receivers should prefer costly signals, as the costs provide a guarantee for honesty. The current results along with previous investigations (see [8] for a review) show that none of the main predictions of the handicap principle are supported. Signals need not be costly to be honest and costlier signals are not necessarily more honest. Therefore, the claim that honest signals *evolve because they are costly* is incorrect. Signals evolve just like any other trait in biology. Efficacy of a functional trait can improve with additional investment, but only up to a point (assuming that it contributes to the potential cost of cheating); after that point, further investment (i.e., waste) is detrimental. To understand whether equilibrium cost plays a role in promoting honesty one has to understand the relationship between equilibrium cost and the potential cost of cheating. The lack of correlation between the two implies that increasing the equilibrium cost of signals will be detrimental to the evolution of honest signalling.

## Founding

S.S. was supported by the National Research, Development and Innovation Office – NKFIH (OTKA) grant K 108974 and by the European Research Council (ERC) under the European Union’s Horizon 2020 research and innovation programme (grant agreement No 648693). D.J.P. acknowledges support from Austrian Science Fund (FWF: P 24711-B21 and P 28141-B25). The funding agencies had no role in the study design, analyses, or publication.

## Author contribution

S.S. and D.J.P conceived the idea, S.S. wrote the IBM and analysed the results, S.S. and D.J.P wrote the paper.

